# Rigid, bivalent CTLA-4 binding to CD80 is required to disrupt the *cis* CD80 / PD-L1 interaction

**DOI:** 10.1101/2024.04.19.590271

**Authors:** Maximillian A Robinson, Alan Kennedy, Carolina T Orozco, Hung-Chang Chen, Erin Waters, Dalisay Giovacchini, Kay Yeung, Lily Filer, Claudia Hinze, Christopher Lloyd, Simon J Dovedi, David M Sansom

## Abstract

The CTLA-4 and PD-1 checkpoints control immune responses to self-antigens and are key targets in cancer immunotherapy. Both pathways are connected via a cis interaction between CD80 and PD-L1, the ligands for CTLA-4 and PD-1 respectively. This cis interaction prevents PD-1 binding to PD-L1 but is reversed by CTLA-4 trans-endocytosis of CD80. However, the mechanism by which CTLA-4 selectively removes CD80 but not PD-L1 is unclear. Here we show that CTLA-4 – CD80 interactions are unimpeded by PD-L1 and that CTLA-4 binding with CD80 does not displace PD-L1 per se. Rather, both the rigidity and bivalency of the WT CTLA-4 molecule is required to orientate CD80 such that PD-L1 interactions are no longer permissible. Moreover, soluble CTLA-4 released PD-L1 only at specific expression levels of CD80 and PD-L1, whereas CTLA-4 trans-endocytosis released PD-L1 in all conditions. These data show that PD-L1 release from CD80 is driven by biophysical factors associated with orientation and bivalent cross-linking of proteins in the cell membrane and that trans-endocytosis of CD80 efficiently promotes PD-L1 availability.

## Introduction

CTLA-4 is a critical immune checkpoint in the attenuation of T-cell responses. Homozygous CTLA-4 deletion in mice leads to a fatal, lympho-proliferative phenotype, a result of excessive T-cell activation due to a deficiency in the regulatory T-cell compartment^1,2^. Homozygous CTLA-4 mutations in humans have not been reported and are assumed fatal; however, patients with heterozygous mutations in the CTLA4 gene often present with severe immune dysregulation and autoimmunity^3^.

CTLA-4 is an endocytic receptor that functions as the primary antagonist of the CD28 co-stimulatory pathway^2^. Both CD28 and CTLA-4 share two ligands CD80 and CD86, however CTLA-4 binds both with a greater affinity than CD28^4^. We have shown CTLA-4 physically depletes CD80 and CD86 in a process called trans-endocytosis^5^. By binding and removing CD80 and CD86 from opposing cells, CTLA-4 can regulate the amount of co-stimulatory ligand available for CD28-driven T-cell activation. Moreover, the identity of the ligand bound also impacts on CTLA-4 fate and the levels of functional CTLA-4^6^. These observations highlight the cell-extrinsic nature of CTLA-4 function, in keeping with experimental results from mouse studies and the observed function of regulatory T-cells^7,8^.

Research into CTLA-4 has yielded several successful clinically approved therapies. Abatacept (CTLA-4-Ig), a CTLA-4-Immunoglobulin (Ig) fusion protein, functions by binding free CD80 and CD86, blocking ligation of CD28 and therefore T-cell responses. Abatacept was first approved for patients with rheumatoid arthritis, but has since been indicated for the treatment of psoriatic arthritis, GVHD and more recently in CTLA-4 deficiency syndromes^9–13^. In contrast, immunotherapies that block CTLA-4 have revolutionised the treatment of cancer and function by increasing CD28 activity, thereby potentiating the T-cell response^14^. However, anti-CTLA-4 therapies have high levels of immune related adverse events which sub-optimally limit pathway inhibition^15^. Moreover, targeting the PD-1 receptor, another cell-surface protein involved in T-cell inhibition, has emerged for many cancer indications. Combination therapies blocking both the CTLA-4 and PD-1 receptors demonstrate increased responses as compared to either CTLA-4 or PD-1 blockade alone^16,17^.

Despite initial concepts that CTLA-4 and PD-1 represent discrete pathways, recently it has emerged that there is a significant molecular overlap between the CTLA-4 and PD-1 pathways. Butte *et al.* first demonstrated that CD80 and PD-L1 interact with a low micromolar affinity^18^; subsequently, CD80 and PD-L1 were shown to predominately interact in *cis*, when both ligands are co-expressed on the same cell membrane^19^. Moreover, the interaction between CD80 and PD-L1 precludes PD-1 binding, thus preventing PD-1 mediated inhibition^19,20^. Mutational analysis suggests that the CD80 and PD-1 binding site on PD-L1 are overlapping, with further structural data confirming that residues on the PD-L1 protein that mediate PD-1 binding are shared with CD80^21^. Thus, it appears likely that CD80 inhibits PD-1 / PD-L1 interactions by steric obstruction of the PD-1 binding site.

In contrast, CD80 remains available for CTLA-4 binding despite being in complex with PD-L1^22^. *In vitro* cellular assays demonstrated that trans-endocytosis of CD80 by CTLA-4 was unimpeded by the CD80 interaction with PD-L1 and that CD80 depletion was remarkably specific, as PD-L1 was retained at the cell membrane despite CD80 removal. Thus, following removal of CD80 by trans-endocytosis, PD-L1 / PD-1 interactions are rescued, and PD-1-mediated inhibition is restored. In principle, this connects CTLA-4 function with the promotion of PD-1 inhibition. Additional reports indicated that soluble CTLA-4 variants (CTLA-4 Ig) and cell-expressed CTLA-4 mutants that are unable to perform trans-endocytosis were also capable of releasing PD-L1, features we did not initially observe in our system^21,23^. This raises several questions about how trans-endocytosis of CD80 occurs without its heterodimeric partner, PD-L1, why the impact of soluble CTLA-4 is variable and how CTLA-4 can disrupt CD80 / PD-L1 interactions.

Herein we explored the mechanism by which CTLA-4 can selectively deplete CD80 but not PD-L1. We observed that soluble CTLA-4 Ig binding to cells expressing both CD80 and PD-L1 only rescues PD-1 binding when both ligands are expressed at similar levels, but not when CD80 was in excess. In addition, we generated a series of soluble CTLA-4 variants to understand why PD-L1 release was so dependent on ligand expression levels. Using both monovalent CTLA-4 constructs and flexible, bivalent CTLA-4 constructs, we demonstrate that neither could cause dissociation of the CD80 / PD-L1 heterodimer. Therefore, release of PD-L1 is not achieved by simple CTLA-4 binding to CD80 / PD-L1 heterodimers *per se* but requires both bivalent and rigid binding of the WT CTLA-4 protein with CD80 to release PD-L1. Finally, we compared the ability of soluble CTLA-4 with cell-expressed WT CTLA-4 for the ability to release PD-L1, and show that, in contrast to soluble CTLA-4, only trans-endocytosis of CD80 could effectively release PD-L1, regardless of CD80 and PD-L1 expression level.

## Results

### CTLA-4 Ig disrupts CD80 / PD-L1 interactions only at defined ligand levels

To study the interaction between CD80 and PD-L1, we transduced into a DG-75 B-cell line GFP-tagged CD80 and mCherry-tagged PD-L1. We sorted on cell populations with varying levels of ligand and determined the molecular ratio of CD80 to PD-L1 in each independent cell line **(Fig 1 A),** based on ligand quantitation we performed previously^22^.

**Fig 1.**
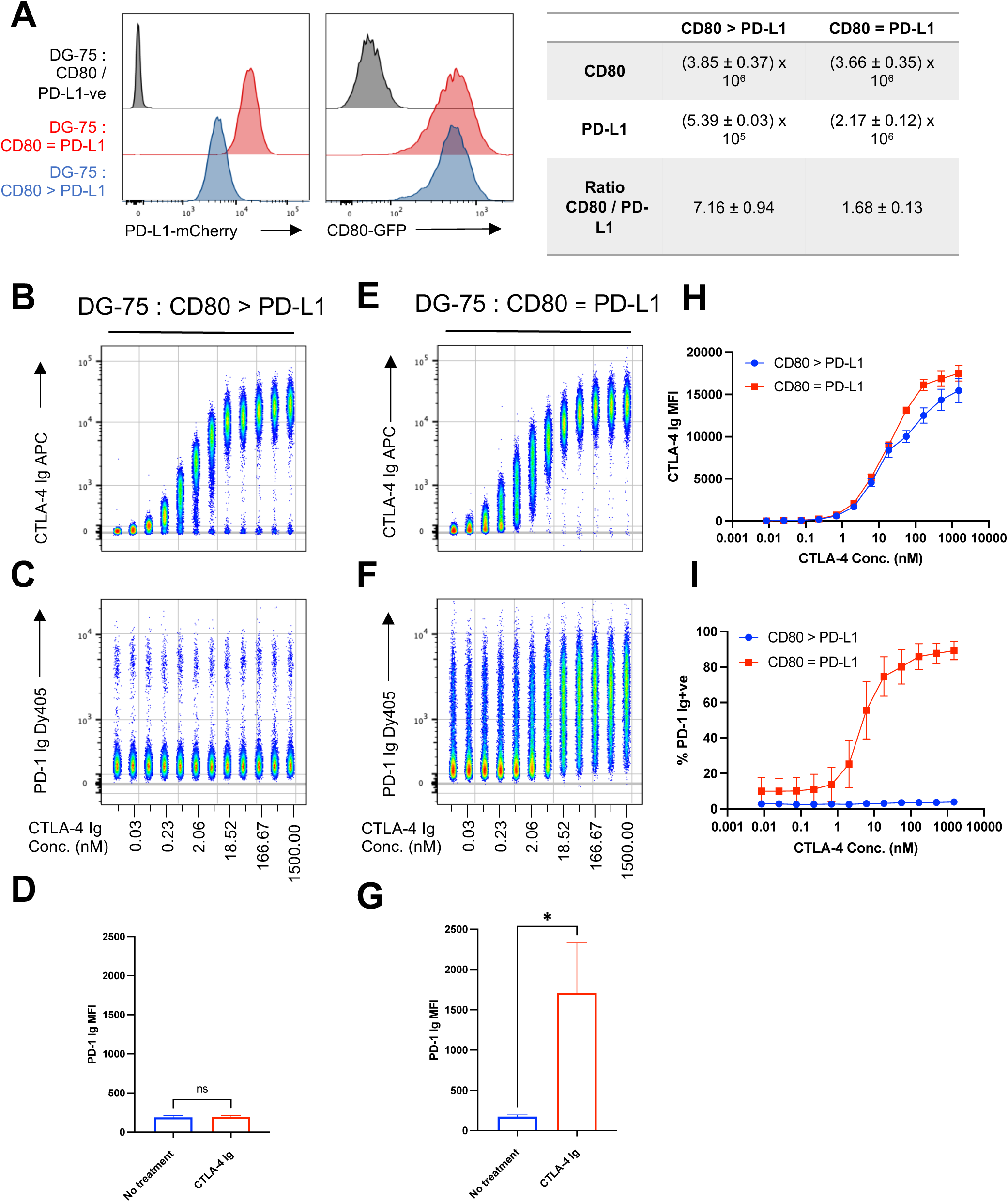
CTLA-4 Ig only disrupts CD80 / PD-L1 interactions at defined ligand levels. **(A)** Representative histograms showing PD-L1-mCherry and CD80-GFP levels on two different DG-75 cell lines. Right-hand table details ligand numbers and molecular ratio of CD80 and PD-L1 on either cell line. **(B)** Concatenated flow cytometry plot of a 12-point serial dilution of CTLA-4 Ig-APC, starting at 1500nM, on DG-75 : CD80 > PD-L1, **(C)** followed by PD-1-Ig-Dy405 detection. **(D)** MFI of PD-1 Ig binding to DG-75 : CD80 > PD-L1 +/-1500 nM of CTLA-4 Ig. **(E - G)** As in (B - D) but performed on DG-75 : CD80 = PD-L1. **(H)** Graphical comparison between DG-75 : CD80 = PD-L1 and DG-75 : CD80 > PD-L1 of CTLA-4 Ig-APC binding, at indicated concentrations. **(I)** Graphical comparison between DG-75 : CD80 = PD-L1 and DG-75 : CD80 > PD-L1 of PD-1 Ig positive cells, after incubation with indicated concentrations of CTLA-4 Ig- APC. Data are representative of three independent experiments showing mean ± SD. *P ≤ 0.05, ns, not significant: paired t-test (D & G).

We then titrated labelled a CTLA-4 Ig fusion protein (CTLA-4 Ig, referred to clinically as Abatacept) on a population of DG-75 cells expressing an excess of CD80 as compared to PD-L1. As observed previously^22^, CTLA-4 Ig binding to CD80 was unimpeded by PD-L1 co-expression (**Fig 1 B**). Next, we used a soluble PD-1 Ig fusion protein (PD-1 Ig) to determine the availability of PD-L1 for PD-1 binding. We observed that when CD80 levels were in excess of PD-L1, PD-1 Ig binding was abrogated, and that this was unaffected by the presence of CTLA-4 Ig (**Fig 1 C**). Even at the highest dose of CTLA-4 Ig, we failed to see a shift in PD-1 Ig binding (**Fig 1 D**). Therefore, CTLA-4 binding to CD80 did not promote PD-L1 release *per se* and a tripartite complex between CD80, PD-L1 and CTLA-4 was permitted.

We then repeated the same assay on DG-75 cells expressing approximately equal levels of PD-L1 as compared to CD80 (DG-75 : CD80 = PD-L1). These cells showed a low level of baseline PD-1 Ig staining, representing some free PD-L1, however this was clearly enhanced at increasing concentrations of CTLA-4 Ig (**Fig 1 E&F**). Moreover, at the highest CTLA-4-Ig doses we observed a significant increase in PD-1 Ig binding (**Fig 1 G**). Importantly, the levels of CTLA-4 Ig binding were comparable between cell lines **(Fig 1H),** yet a concomitant increase in PD-1 Ig binding was only observed on the DG-75 : CD80 = PD-L1 population **(Fig 1I).** Thus, we concluded that soluble CTLA-4 can enhance PD-L1 / PD-1 binding, although only at favourable ratios of CD80 to PD-L1 expression.

CD28 shares significant similarities with CTLA-4: both are covalent homodimers and share the MYPPPY motif, essential for CD80 binding^24^. We therefore tested the impact of CD28 binding on PD-L1 release. We produced a soluble CD28 Ig fusion protein (CD28 Ig) and, using biolayer interferometry (BLI), confirmed immobilised CD28 Ig bound monomeric His tagged-CD80 (**Fig S1 A**) with a weaker affinity (CD28 / CD80 K_D_ = (4.27 ± 0.46) × 10^−5^) than CTLA-4, as expected^25^. Furthermore, when titrated onto DG-75 cells expressing equal levels of CD80 and PD-L1, CD28 Ig binding to CD80 was “right-shifted” as compared to CTLA-4 Ig, reflecting its lower affinity (**Fig S1 B**). Intriguingly, despite CD28 sharing the same binding site as CTLA-4, we failed to observe any release of PD-L1, as measured by PD-1 Ig binding (**Fig S1 C**). Considering the binding site on CD80 is shared between CTLA-4 and CD28, these results suggest that CTLA-4 binding to CD80 is uniquely capable of releasing PD-L1.

### CTLA-4 monovalent binding fails to disrupt CD80 / PD-L1 interactions

Despite CD28 and CD28 Ig existing as covalent homodimers, structural data suggests that CD28 / CD80 interactions are primarily monovalent, due to the angle of ligand binding preventing both CD28 sites being engaged simultaneously^26^. In contrast, CTLA-4 can interact with CD80 bivalently^27^. We therefore considered the hypothesis that bivalent binding may be required to drive the observed dissociation of CD80 from PD-L1 following CTLA-4 binding.

To test this, we generated a soluble, monovalent CTLA-4 Ig fusion protein (mono-CTLA-4 Ig) using the knobs-into-holes technology (see methods and **Fig S2** for protein sequences)^28^. Affinity measurements indicated mono-CTLA-4 Ig bound CD80 with a similar K_D_ to WT CTLA-4 Ig, suggesting that CTLA-4 / CD80 affinity was essentially the same for both constructs (**Fig 2 A & B**). To confirm our mono-CTLA-4 Ig construct bound monovalently, we performed a BLI assay using immobilised CD80 His, probing a range of concentrations of WT CTLA-4 Ig and mono-CTLA-4 Ig (**Fig S3 A**). Even at low concentrations, WT CTLA-4 Ig failed to dissociate from CD80, a result of the enhanced avidity of the bivalent interaction (**Fig S3 B**). In contrast, mono-CTLA-4 Ig could readily dissociate from immobilised CD80, consistent with a monovalent binding model between the ligand-receptor pair (**Fig S3 C**). DG-75 : CD80 = PD-L1 cells were then treated with a titration of mono-CTLA-4 Ig (**Fig 2 C**). Remarkably, even at saturating doses of mono-CTLA-4 Ig, we were unable to restore PD-1 Ig binding (**Fig 2 C&D**). Taken together, these results show that CTLA-4 / CD80 binding alone is insufficient to release PD-L1 but that CTLA-4 mediated dissociation of the CD80 / PD-L1 heterodimer is dependent upon bivalent CTLA-4 binding.

**Fig 2.**
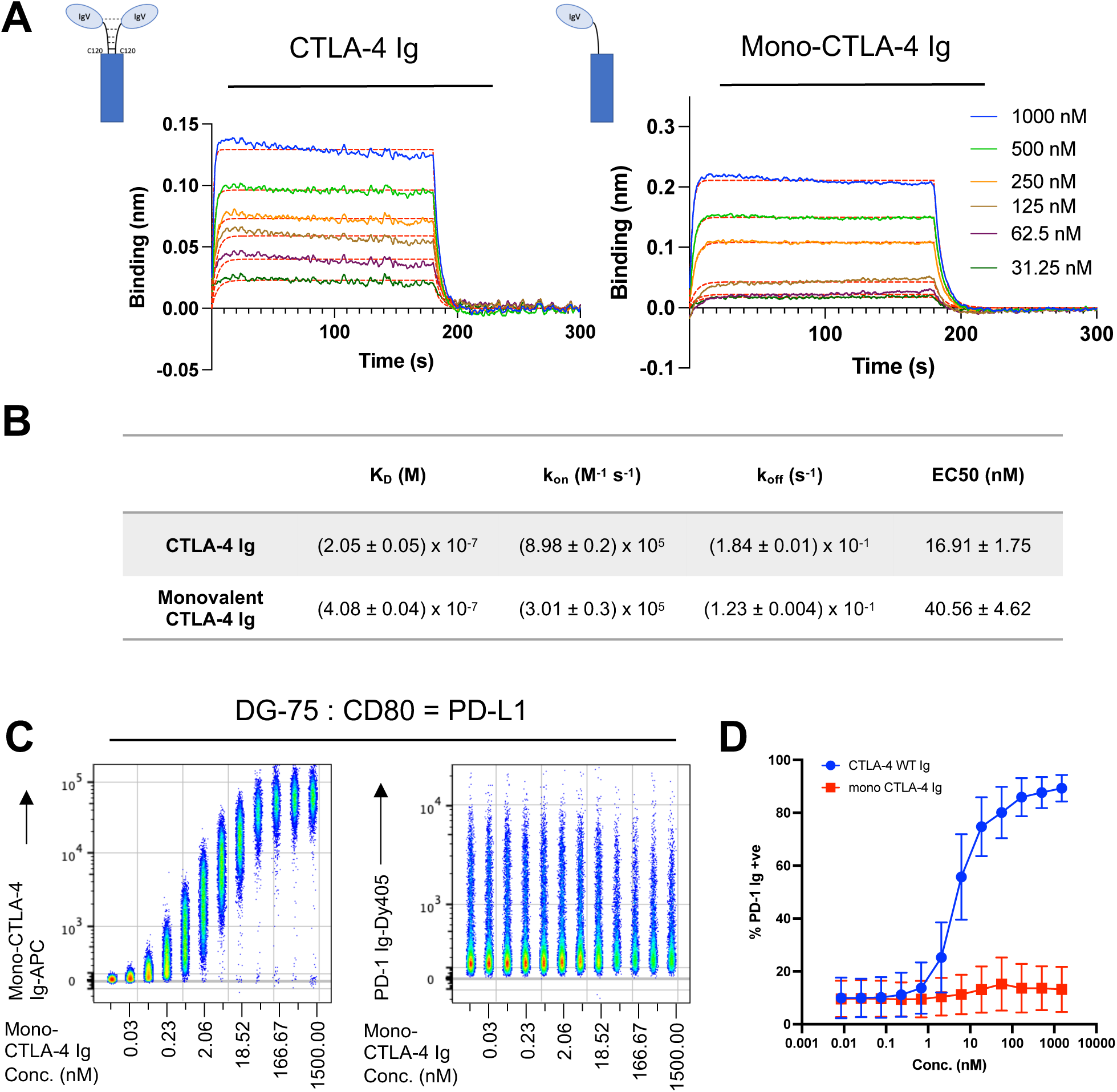
CTLA-4 monovalent binding fails to disrupt the CD80 / PD-L1 interaction. **(A)** Binding curves showing the association and dissociation of 1000, 500, 250, 125, 62.5, 31.3 nM of CD80-His to CTLA-4 Ig (left) and monovalent CTLA-4 Ig (right). Red lines show best fit to a 1:1 binding model. **(B)** Kinetic and thermodynamic parameters (K_D_, k_on_, k_off_) obtained from the best global fit of the association/dissociation data to a 1:1 binding model. The errors given are fitting errors from the global fitting. **(C)** Concatenated flow cytometry plot of a 12-point serial dilution of mono-CTLA-4 Ig-APC, starting at 1500 nM, on DG-75 : CD80 = PD-L1, followed by PD-1 Ig-Dy405 detection (right-hand panel). **(D)** Graphical representation of PD-1 Ig positive cells after treatment with WT CTLA-4 Ig or monovalent CTLA-4 Ig, at indicated concentrations. Data are representative of three independent experiments showing mean ± SD.

### Structural modelling of CTLA-4 interacting with the PD-L1 / CD80 heterodimer

CD80 is composed of a membrane distal immunoglobulin variable (IgV) domain and a membrane proximal immunoglobulin constant (IgC1) domain, which forms transient, non-covalent homodimers (mediated via IgV domain interactions). These dimers interact with low affinity and a reported K_D_ of 20-50 □M^29^. CD80 dimerisation results in two monomers arrayed parallel to one another, and orthogonal to the cell membrane in a conformation that is maintained when complexed with CTLA-4 **(Fig 3A)**^27^.

**Fig 3.**
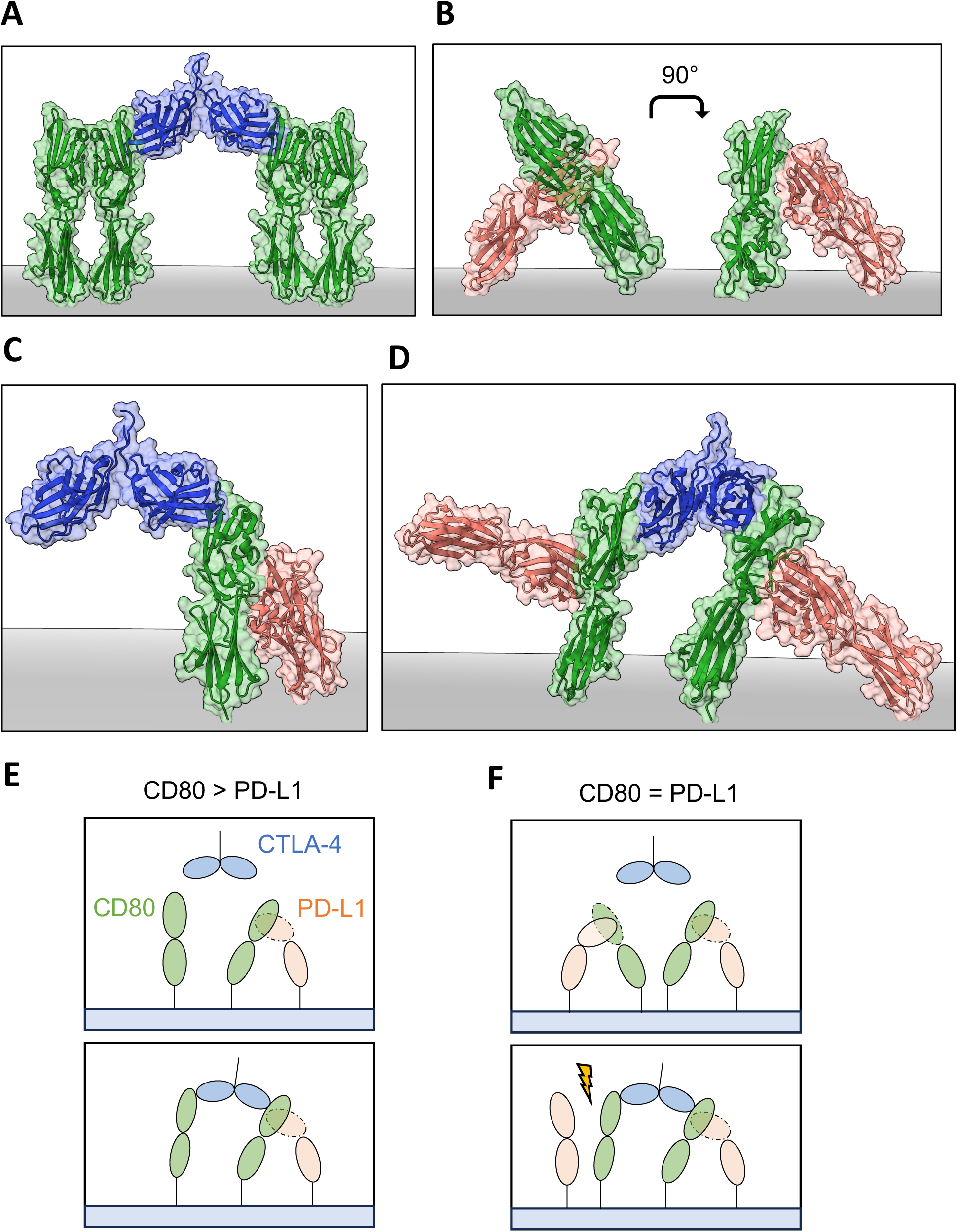
Structural models of CTLA-4 interacting with the PD-L1/CD80 heterodimer. **(A)** Representative surface and ribbon structure of CTLA-4 (blue) in complex with two CD80 homodimers (green). (PDB: 1I8L) **(B)** Rotated view of the ALPN-202 CD80 vIgD / PD-L1 ECD (red) asymmetric unit aligned with wild type CD80 (green) (PDB: 7TPS and 1DR9). **(C)** Model alignments of the CD80 / PD-L1 complex interacting with CTLA-4 monovalently. **(D)** Bivalent CTLA-4 binding to PD-L1/CD80 heterodimers (PDB: 7TPS, 1DR9 and 1I8L); structure has been rotated to highlight the angle of CD80 relative to the membrane and displacement of the PD-L1 protein. **(E & F)** Schematic depicting CTLA-4 (blue) binding to CD80 (green) / PD-L1 (red) heterodimers when CD80 is expressed in excess to PD-L1 (E) and when CD80 and PD-L1 are expressed at equimolar ratios (F).

Although the full crystal structure of CD80 interacting with PD-L1 is yet to be determined, a recent paper resolved the structure of a variant high affinity CD80 IgV domain (ALPN-202 CD80 vIgD) in complex with PD-L1^30^. We therefore aligned the structure of WT CD80 based upon the high affinity CD80 / PD-L1 crystal **(Fig 3B)**. In this model PD-L1 binds the CD80 dimer interface at an unusual angle (“lying down”) such that it would appear to prevent CD80 from existing at an orthogonal angle with the membrane. Nonetheless, the model indicates that both CTLA-4 and PD-L1 can simultaneously bind to CD80 **(Fig. 3C)**. However, by changing the angle of CD80 in the membrane, CTLA-4 interactions are then skewed when compared to CD80 homodimer binding. Furthermore, it appears unlikely that CTLA-4 can bridge two CD80 / PD-L1 complexes, since both PD-L1 molecules could no longer anchor in the membrane **(Fig 3D).** Structural analysis therefore suggests that bivalent binding of CTLA-4 to two CD80 / PD-L1 heterodimers is unlikely, offering an explanation as to why release of PD-L1 depends on the relative levels of CD80 and PD-L1 in the membrane.

Accordingly, if CD80 levels are in excess to PD-L1, membrane CD80 will form a mixed population of transient CD80 homodimers and CD80 / PD-L1 heterodimers. Thus, CTLA-4 could bivalently bind with one arm to a CD80 / PD-L1 heterodimer, with the other arm binding a CD80 monomer or homodimer **(Fig 3E)**. In the context of equimolar expression CD80 and PD-L1, the large majority of CD80 ligand is complexed with PD-L1. Here, bivalent binding is precluded until CD80 and PD-L1 dissociate, only then allowing CTLA-4 to bridge a PD-L1 / CD80 heterodimer and a CD80 monomer **(Fig 3F).** Moreover, once formed, bivalent CTLA-4 / CD80 complexes would likely prevent PD-L1 re-association and therefore generate detectable free PD-L1. Therefore, the only condition in which we observe significant, but ultimately partial, restoration of PD-1 / PD-L1 binding by soluble CTLA-4 is under conditions where the vast majority of CD80 at the membrane is in complex with PD-L1.

### Rigid body CTLA-4 binding is required to restore PD-1 binding

The above model suggests that, in addition to bivalency the geometry of CTLA-4 interacting with CD80 is important. This geometry is constrained both by the binding epitope but also by the rigidity of the CTLA-4 dimer. Indeed, the CTLA-4 / CD80 interaction has been described as rigid body, since neither molecule’s conformation is changed when comparing their *apo* and complexed crystal structures^31^. We therefore tested whether CTLA-4 rigid body binding is also required to mediate PD-L1 release by producing flexible CTLA-4 molecules. We produced two CTLA-4 Ig variants: one where we substituted the native cysteine, required for the disulfide bridge, with an alanine (CTLA-4 C120A Ig); and a second variant where we removed the CTLA-4 stalk region and replaced with a flexible GGGGS linker (CTLA-4 V114 G4S Ig) (**Fig S2**).

BLI analysis confirmed these constructs were both able to bind CD80. Immobilised CTLA-4 C120A Ig and CTLA-4 V114 G4S Ig bound soluble CD80-His with a similar affinity to WT CTLA-4 Ig (**Fig 4 A-B**); given that binding between CTLA-4 and CD80 utilises a ligand binding site in the IgV domain, alterations in the stalk region had little impact on binding, as expected. Using these constructs in a BLI assay with immobilised CD80-His, we observed slow dissociation (as seen with WT CTLA-4 Ig), indicating avidity enhanced bivalent interactions still occurred with these constructs (**Fig S3 D-E**).

**Fig 4.**
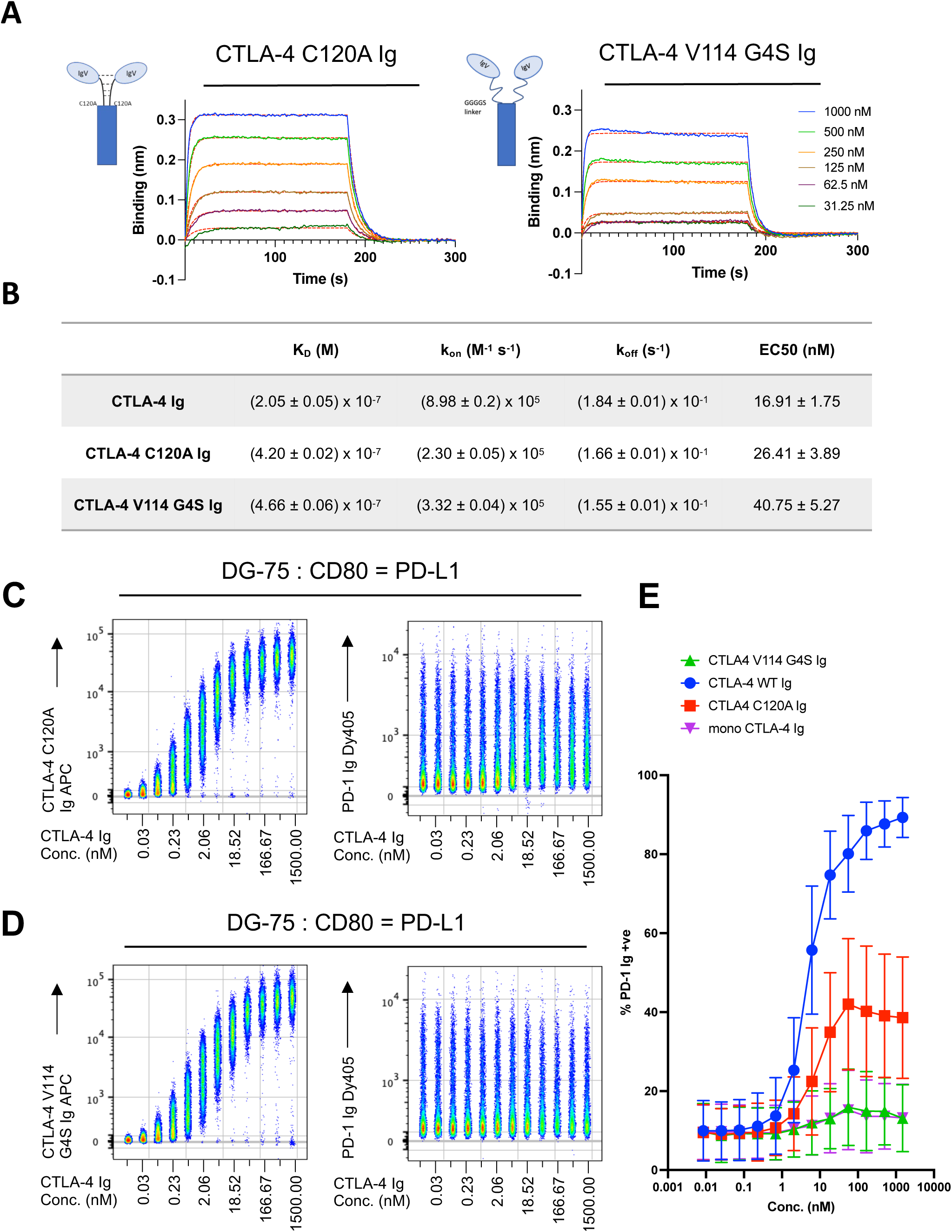
Rigid body CTLA-4 binding is required to restore PD-1 binding. **(A)** Binding curves showing the association and dissociation of 1000, 500, 250, 125, 62.5, 31.3 nM of CD80-His to CTLA-4 C120A Ig (left) and CTLA-4 V114 G4S Ig (right). Red lines show best fit to a 1:1 binding model. **(B)** Kinetic and thermodynamic parameters (K_D_, k_on_, k_off_) obtained from the best global fit of the association/dissociation data to a 1:1 binding model. The errors given are fitting errors from the global fitting. **(C & D)** Concatenated flow cytometry plot of a 12-point serial dilution of CTLA-4 C120A Ig-APC (C), and CTLA-4 V114 G4S Ig-APC (D), starting at 1500 nM, on DG-75 : CD80 = PD-L1, followed by PD-1-Ig-Dy405 detection (right-hand panels). **(E)** Graphical representation of PD-1 Ig positive DG-75 : CD80 = PD-L1 cells, after treatment with indicated CTLA-4 Ig construct, at indicated concentrations. Data are representative of three independent experiments showing mean ± SD.

When these flexible, bivalent constructs were titrated on to DG-75 : CD80 = PD-L1, we observed a similar EC50 to WT CTLA-4 Ig, demonstrating that our CTLA-4 Ig variants readily bound CD80 in complex with PD-L1 (**Fig 4 B**). However, at saturating concentrations, both CTLA-4 C120A Ig and CTLA-4 V114 G4S Ig failed to restore PD-1 Ig binding to the level seen with WT CTLA-4 Ig (**Fig 4 C-D**). Interestingly, the highly flexible CTLA-4 V114 G4S Ig failed to alter PD-1 Ig binding whatsoever, indicating that structural rigidity was required to dissociate the CD80 / PD-L1 heterodimer. However, we observed that, at high doses, CTLA-4 C120A Ig could release low levels of PD-L1, as measured by PD-1 Ig binding, suggesting an intermediate phenotype (**Fig 4 E**). Therefore, comparing three novel constructs (mono-CTLA-4 Ig, CTLA-4 C120A Ig and CTLA-4 V114 G4S Ig) we concluded that the ability to restore PD-1 Ig binding and effect PD-L1 release required rigid, bivalent CD80-CTLA-4 interactions.

### Native cell-expressed CTLA-4 is uniquely capable of releasing PD-L1 regardless of CD80 expression levels

Finally, we compared the ability of soluble CTLA-4-Ig proteins with cell expressed WT CTLA-4 to release PD-L1. Here we incubated DG-75 cells co-expressing CD80 and PD-L1 with either a WT CTLA-4+ve Jurkat T-cell, or with a CTLA-4-negative Jurkat supplemented with 50nM of CTLA-4 Ig, a dose sufficient to saturate CD80. Under equimolar conditions of PD-L1 and CD80 expression **(Fig 5A)** both CTLA-4 Ig and cell-expressed WT CTLA-4 were able to release PD-L1, as measured by PD-1 Ig binding. When comparing our generated CTLA-4 variants, only CTLA-4 C120A Ig was able to partially release PD-L1, although at much lower levels than WT CTLA-4 Ig **(Fig S4)**. Interestingly, CTLA-4-Del36, a CTLA-4 mutant lacking the cytoplasmic tail essential for trans-endocytosis, also partially restored PD-1 binding but proved much less effective than CTLA-4 Ig or cell-expressed WT CTLA-4. In contrast, when using DG-75 cells expressing excess CD80, which is a more exacting scenario, only cell-expressed WT CTLA-4 was capable of restoring PD-1 Ig binding **(Fig 5B)**. Moreover, cell-expressed WT CTLA-4 was able to deplete CD80 ligand by trans-endocytosis, as measured by loss of CD80-GFP from the DG-75 **(Fig 5C&D)**.

**Fig 5.**
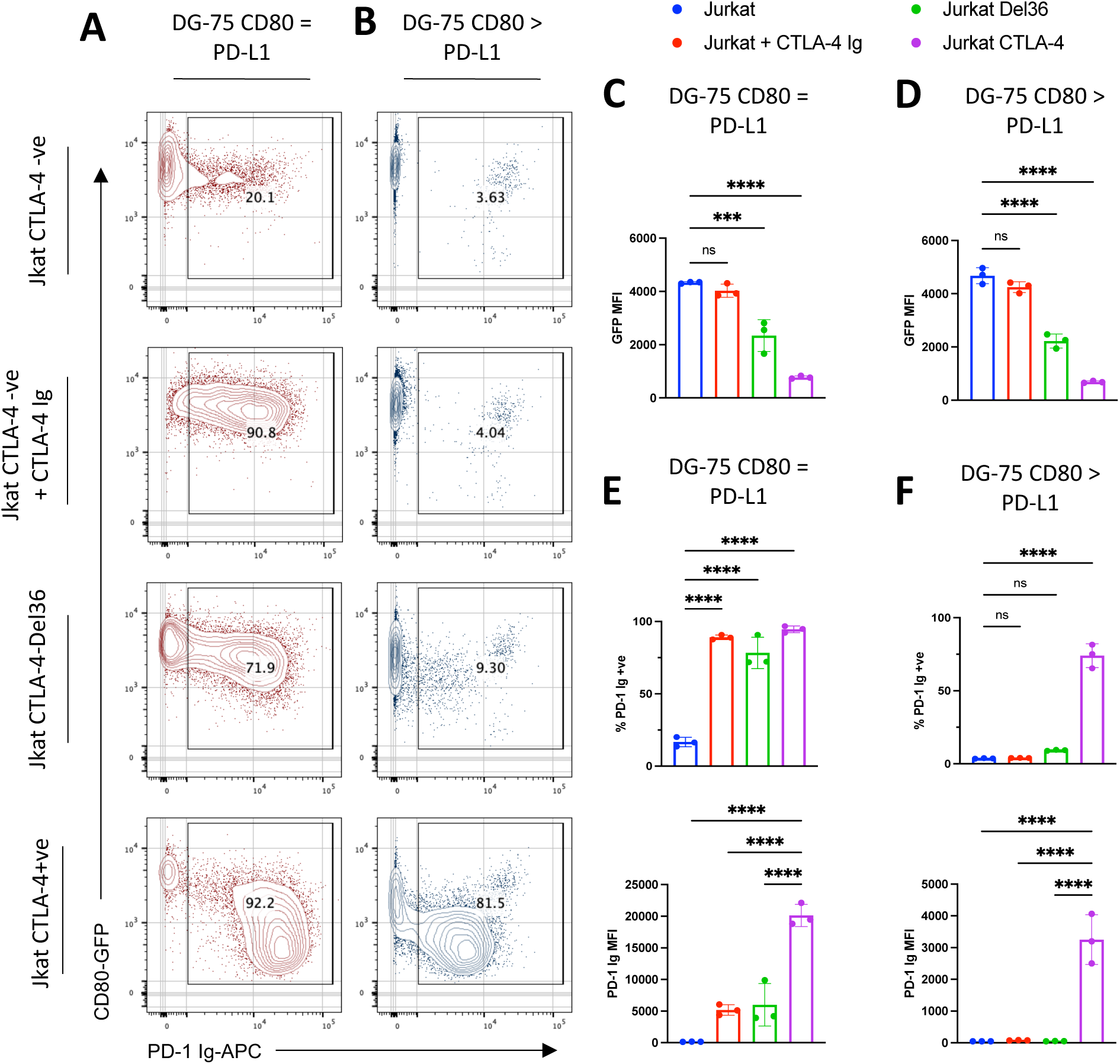
Only cell-expressed CTLA-4 restores PD-1 binding irrespective of CD80 and PD-L1 expression levels. **(A & B)** Contour plots showing DG-75 : CD80 = PD-L1 (A) and DG-75 : CD80 > PD-L1 (B) incubated for 24hrs with: CTLA-4-ve Jurkat, CTLA-4-ve Jurkat + 50nM CTLA-4 Ig, CTLA-4-Del36 Jurkat, CTLA-4+ve Jurkat. Cells were stained with 1ug/ml of PD-1 Ig-APC after incubation. Data shows representative FACS plots of CD80-GFP vs. PD-1 Ig. **(C-F)** Graphical representation of (A & B) respectively, plotting CD80-GFP MFI, % of PD-1 Ig +ve cells and PD-1 Ig MFI on DG-75 : CD80 = PD-L1 (C,E) and DG-75 : CD80 > PD-L1 (D,F). Data are representative of three independent experiments showing mean ± SD. ***P ≤ 0.001, ****P ≤ 0.0001, ns, not significant: one-way ANOVA with Tukey’s multiple comparisons test (C-F).

In addition, although soluble CTLA-4 restored PD-1 Ig binding to the entire DG-75 population expressing equimolar levels of CD80 and PD-L1 **(Fig 5E, top graph)**, incubation with WT CTLA-4+ve Jurkats resulted in increased PD-1 Ig binding, as measured by MFI **(Fig 5E, bottom graph)**. Here we observed an almost four-fold difference in PD-1 Ig MFI between CTLA-4 Ig release of PD-L1 and cell-expressed WT CTLA-4, suggesting cell-bound CTLA-4 is a more effective at releasing PD-L1 when compared with soluble CTLA-4 Ig and CTLA-4 Del36. These results highlight the importance of endocytic, cell-expressed CTLA-4, which is capable of both binding to CD80 and physically depleting the ligand. In contrast, CTLA-4 Ig and CTLA-4-Del36 are less able to release PD-L1 and are dependent on permissive levels of CD80 and PD-L1 expression **(Fig 5E vs 5F)**. Thus, only through trans-endocytosis of CD80 can CTLA-4 effectively release total PD-L1, regardless of expression levels of CD80 to PD-L1.

## Discussion

The CTLA-4 and PD-1 pathways are major targets for therapeutic intervention in the treatment of autoimmunity, transplantation, and cancer. Recently a significant molecular overlap between these pathways has been identified, mediated by the *cis* CD80 / PD-L1 interaction. The impact of this interaction is to disable PD-L1 function and to place it under the control of CD80 and CTLA-4 in some circumstances.

Several studies have now shown that by forming heterodimers with PD-L1, CD80 physically obstructs the PD-1 binding site, thereby acting as an antagonist to PD-1 signalling^19,20^. The impact on CD28 and CTLA-4 appears much more subtle, consistent with the CD28 / CTLA-4 binding site being on the opposite face of CD80 as compared with the PD-L1 binding interface. Whilst one study suggesting that CD80 / PD-L1 interactions affected CTLA-4 function^32^, these results contrast to several others where the impact on CTLA-4 binding appears intact ^21–23^.

We previously demonstrated that cell-expressed CTLA-4 could both bind and trans-endocytose CD80, despite its interaction with PD-L1^22^. Interestingly, depletion of CD80 by CTLA-4 demonstrated the remarkable specificity of trans-endocytosis, as removal of CD80 had no impact on PD-L1 levels, despite clear evidence of CD80 / PD-L1 heterodimers on the membrane. Thus, in the context of CD80 and PD-L1 co-expression, CTLA-4 trans-endocytosis of CD80 restores competent PD-L1 at the membrane and thereby promotes PD-1-mediated inhibition. Consistent with our findings, studies by Tekguc *et al.* likewise found that soluble CTLA-4 interactions and CTLA-4 trogocytosis could also liberate PD-L1^23^. However, the mechanism by which CTLA-4 liberates free PD-L1 remains unresolved. Here we have investigated the requirements for CTLA-4 to liberate PD-L1 from CD80 / PD-L1 heterodimers found in membranes. Our data reveal a mechanism whereby rigid, bivalent binding of CTLA-4 is required to orientate CD80 in the membrane such that PD-L1 binding is no longer favourable. For soluble bivalent CTLA-4 molecules and CTLA-4 molecules incapable of trans-endocytosis, partial release of PD-L1 only occurs at specific (approximately equimolar) ratios of CD80 to PD-L1. In contrast, cell-expressed WT CTLA-4 molecules can utilise trans-endocytosis to continually deplete CD80, allowing PD-L1 release regardless of CD80 / PD-L1 expression ratios.

Several lines of evidence support the above conclusions. A recent crystal structure of a high-affinity CD80 vIgD domain in complex with human PD-L1^30^ has provided new insights. Here, Maurer *et al.*, showed that the CD80 binding interface for PD-L1 is situated along the (ABED) strands of the membrane distal IgV domain, in line with previous mutational data^20,21^. CD80 homodimerization, which also uses this interface^29^, is therefore incompatible with PD-L1 binding. In membranes, CD80 can form CD80 homodimers or heterodimers with PD-L1, but may also present as monomers due to the low affinity and non-covalent nature of both the homodimer and heterodimer^18,29,33^. These structural data further highlight that CTLA-4 binding to the opposite (GFCC’C’’) face of the CD80 IgV domain is therefore unimpeded by PD-L1. Thus, there appears no obvious structural incompatibility between CTLA-4 and PD-L1 binding to the same CD80 molecule and forming a tripartite complex. Moreover, there is no obvious reason why CTLA-4 binding to CD80 should physically displace PD-L1. Indeed, this is compatible with our observations using soluble monovalent or flexible CTLA-4 molecules, which bind to CD80 / PD-L1 heterodimers but do not trigger release of PD-L1 under any circumstances.

Using the alignment of the vIgD CD80 / PD-L1 and WT CD80 structures, it becomes clear that the PD-L1 / CD80 interaction occurs with PD-L1 adopting an unusual “lying down” position. In homodimers, CD80 molecules are essentially parallel to each other and orthogonal to the membrane. However, Maurer et.al,^30^ commented that the orthogonal angle of CD80 in the membrane would be unfavourable for CD80 / PD-L1 heterodimer formation. Accordingly, in the CD80 / PD-L1 heterodimer, CD80 can no longer maintain a 90-degree angle required to form the ‘upright’ CD80 homodimer. Interestingly, previous reports describing the structure of PD-1 – PD-L1 interactions highlighted the remarkable flexibility of the PD-L1 protein^34^, which may accommodate the ‘lying down’ orientation. Controlling the angle of CD80 in the membrane therefore appears crucial to the ability of CTLA-4 to disrupt the PD-L1/CD80 heterodimer. Our observations suggest that bivalent CTLA-4 binding to two CD80 / PD-L1 heterodimers is precluded. However, due to the relatively weak CD80 / PD-L1 heterodimer affinity^18,33^, CD80 and PD-L1 will associate and dissociate naturally, raising the possibility that binding of CD80 to CTLA-4 during dissociation would force CD80 upright in the membrane, thus preventing its re-association with PD-L1.

Prevention of CD80 re-binding to PD-L1 appears to be dependent on the rigidity of CTLA-4. CTLA-4 is normally expressed as a covalent homodimer by virtue of a disulphide bond located at the base of the stalk region. The stalk is further formed of reciprocal hydrophobic and hydrophilic interactions up to the A’ strand of both IgV domains^31^. Such non-covalent interactions, supported by the disulfide bridge at the base of the homodimer, lend remarkable rigidity to the fixed angle between the two IgV domains of the CTLA-4 homodimer. It is interesting to note that CD28 has a different dimer angle, which does not allow bivalent CD80 binding^26^, consistent with its inability to release PD-L1. We observed that replacing the CTLA-4 stalk region with a flexible Glycine-Serine linker removed the ability of CTLA-4 to promote dissociation of the PD-L1/CD80 heterodimer. Thus, a flexible CTLA-4 dimer can bind bivalently to CD80 / PD-L1 complexes without releasing PD-L1. Interestingly, when only the cysteine residue is mutated to an alanine, and therefore retaining only the non-covalent interactions in the stalk region, CTLA-4 C120A Ig appeared to partially release PD-L1, although at much lower levels than WT CTLA-4 Ig. Thus, our data argue that both bivalent and rigid binding, associated with the natural CTLA-4 homodimer configuration, is required to orient CD80 in an upright fashion in the membrane, thereby precluding CD80 / PD-L1 interactions.

We also observed that rigid-bivalent CTLA-4 Ig is only able to release PD-L1 when CD80 and PD-L1 are expressed at approximately equimolar ratios, yet incapable of disrupting the CD80 / PD-L1 interaction when CD80 ligand is in excess of PD-L1. Under conditions of excess CD80 expression, CD80 homodimers will be common and CTLA-4 Ig will bridge two heterodimers infrequently. We hypothesise that flexibility of monomeric (and potentially homodimeric) CD80 is such that CTLA-4 bivalent binding between a CD80 / PD-L1 heterodimer and CD80 is not precluded **(Fig 6A),** thereby maintaining the CD80 / PD-L1 interactions and not releasing PD-L1. In contrast, only where crosslinking of two CD80 / PD-L1 heterodimers is likely, then CTLA-4 Ig promotes the release of PD-L1 **(Fig 6B)**. Crucially, depletion of CD80 by CTLA-4-mediated trans-endocytosis continually adjusts the ratio of CD80 to PD-L1 at the cell membrane. As CD80 is depleted the relative frequency of CD80 / PD-L1 heterodimers increases, eventually reaching equimolar ratios of both ligands, when CTLA-4 bivalency and rigidity can also contribute to the disruption of CD80 / PD-L1 heterodimers. This ability to continually remove CD80 by cell-expressed CTLA-4 explains why WT CTLA-4 is capable of potentiating PD-L1 – PD-1 interactions regardless of CD80 expression levels.

**Fig 6.**
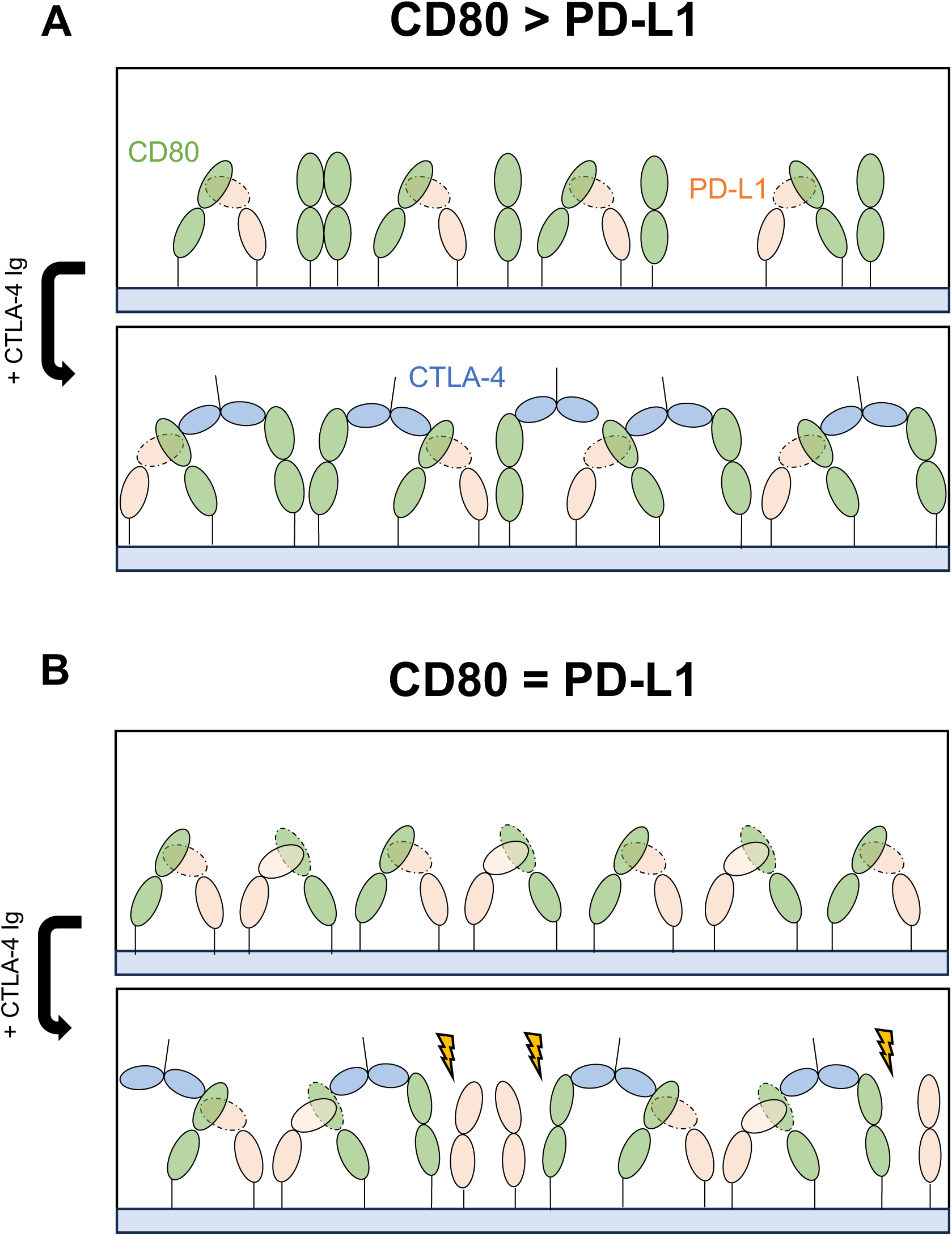
Soluble CTLA-4 binding requires specific CD80 and PD-L1 expression levels to release PD-L1. **(A)** Upon excess CD80 (green) expression as compared to PD-L1 (red), CD80 will present at the membrane as a mixed population of hetero- and homodimers. Upon CTLA-4 Ig (blue) treatment (bottom panel), CTLA-4 Ig can bivalently bind to CD80 and a CD80/PD-L1 complex, without releasing PD-L1. **(B)** Upon equal expression of CD80 and PD-L1, CD80 will exclusively form heterodimers at the membrane. Upon CTLA-4 Ig treatment (bottom panel), CTLA-4 Ig will bivalently bind to two CD80/PD-L1 complexes, which it is unable to maintain due to the angle of CD80 to the membrane: in this case we see partial release of PD-L1 (golden bolts).

The above concepts are relevant to Abatacept, a dimeric CTLA-4 Ig fusion protein that has been clinically approved for autoimmunity^9–11,13^. One possibility is that Abatacept could function to some extent via promoting PD-L1 - PD-1 interactions. However, our data would suggest that this is unlikely for two reasons. First, only specific ligand expression levels are permissive for disruption of the CD80 / PD-L1 interactions, meaning that this effect would be limited to certain APC populations. Second, the MFI of PD-1 Ig binding after treatment with CTLA-4 Ig suggests that CTLA-4 Ig may only partially release PD-L1, in contrast with cell-expressed WT CTLA-4, which can potentially liberate all PD-L1. Moreover, abatacept may prevent cell-expressed CTLA-4 from interacting with CD80 and thereby preclude release of PD-L1 via trans-endocytosis. Another relevant consideration is that a secreted soluble CTLA-4 isoform might also affect PD-L1 activity^35,36^. However, this splice variant lacks the cysteine responsible for forming covalent dimers and displays changes in the stalk region which appear to preclude both non-covalent and covalent interactions. Soluble CTLA-4 is therefore likely to be monomeric and unable to release PD-L1.

Taken together, our data provides an explanation for the specific removal of CD80 and release of PD-L1 by CTLA-4. Our observations indicate that structural characteristics, including bivalency and rigidity of CTLA-4, alongside the angle of CD80 in complex with PD-L1 relative to the cell membrane, are essential for disruption of the CD80 /

PD-L1 heterodimer. Moreover, our results indicate that release of PD-L1 by soluble CTLA-4 is determined by levels of CD80 expression, whereby PD-L1 may only be released after sufficient depletion of CD80. Our previous studies indicated that CD80 trans-endocytosis can also drive CTLA-4 ubiquitination^6^; how CD80 / PD-L1 heterodimers influence this aspect of CTLA-4 biology remains to be determined. Taken together, data herein shows how CD80, PD-L1 and CTLA-4 act in concert to influence PD-1 biology, suggesting that CD80 can act as a master switch influencing the behaviour of both the CTLA-4 and PD-1 pathways.

### Materials and Methods Cell culture

Jurkat E6.1T cells (ATCC, TIB-152) and DG-75 B cells (ATCC, CRL-2625) were cultured in RPMI 1640 (Gibco) supplemented with 10% FCS (Sigma), 2 mM L-Glutamine, 100 U/ml penicillin and 100 mg/ml streptomycin (Gibco). Cell lines were kept at 37°C, 5% CO_2_ in a humidified atmosphere.

Endogenous CD80 was KO with the CRISPR/Cas9 system from the DG-75, Burkitt’s’ B-cell lymphoma cell line. DG-75 cells were then transduced with CD80 and PD-L1 tagged at their C-terminus with GFP and mCherry respectively. Jurkat T-cells were transduced with WT CTLA-4 or CTLA-4 Del 36 cDNAs, the latter generated by insertion of a stop codon that interrupts transcription of the final 36 amino acids of the CTLA-4 cytosolic tail.

Cell line engineering was carried out as described previously^22^.

### Generation of CTLA-4 Ig fusion constructs

For this study, we generated three CTLA-4 hIgG1 Fc fusion constructs, with the peptide sequences of the CTLA-4 ectodomain specified in Figure S2. The CD28 Ig fusion protein included the full ectodomain of CD28 (N19-P152) fused to a hIgG1 Fc domain. The Fc domain used in these constructs contains a triple mutation (TM) which reduces Fc-mediated binding^37^.

To generate our constructs, DNA sequences of the various CTLA-4 and CD28 ectodomains and human Fc TM, knob Fc TM and hole Fc TM were codon-optimised for Chinese Hamster Ovarian (CHO) expression, ordered on GeneArt, and cloned into a proprietary mammalian expression vector with the NEBuilder® HiFi DNA Assembly kit (New England Biolabs). Monovalent CTLA-4 constructs were generated using the knobs-into-holes technology^38^, whereby the pairing of two distinct heavy chains is achieved by molecular complementarity of separate ‘knob’ and ‘hole’ heavy chains. Here, the sequence for CTLA-4 with the C120A mutation was inserted into a human gamma-1 constant heavy chain carrying the ‘knob’ mutation. The ‘knob’ heavy chain was paired with an empty ‘hole’ heavy chain, thus resulting in an Ig construct containing a single CTLA-4 monomer. Plasmid DNA was purified (QIAprep Spin Miniprep Kit, Qiagen), and successful cloning confirmed by Sanger sequencing (GENEWIZ, Azenta). The proteins were transiently expressed in CHO cells using proprietary medium^39^.

6 days post-transfection cells were pelleted, and supernatant vacuum filtered. Cleared supernatant was loaded onto a MabSelect SuRe column (Cytiva) pre-equilibrated with DPBS (Dulbecco’s Phosphate Buffer Saline, Gibco) pH 7.4. Proteins were eluted with 0.1 M glycine pH 2.7 and immediately injected onto a size exclusion chromatography column (SEC, Superdex 200 Increase HiScale 26/40, Cytiva) equilibrated and ran in DPBS pH 7.4. Purity and chain composition were assessed by analytical size exclusion chromatography, with a purity threshold >85% of monomeric protein (Agilent HPLC (1260 Infinity II LC System), TSKgel® G2000SWXL HPLC Column, phase diol, L × I.D. 30 cm × 7.8 mm, 5 μm particle size).

### BLI analysis

The affinity of CTLA-4 and CD28 constructs for CD80 was measured by BLI with the OctetRED384 (Sartorius). All samples were prepared in BLI buffer (DPBS supplemented with 0.02% Tween 20 (Sigma-Aldrich) and 0.1% bovine serum albumin (BSA, Sigma-Aldrich)). Serial dilutions of Human CD80, His Tag (CD80-His, Acrobiosystem, cat # B71-H5228) were made at 1000, 500, 250, 125, 62.5, 31.25, 0 nM. For CD28 kinetic measurements, serial dilution of CD80-His was made at 20, 10, 5, 2.5, 1.25, 0.625 uM.

Biosensors were equilibrated in BLI buffer for 10 mins before start of assay. Baseline measurements were taken before loading of CTLA-4 or CD28 construct on Anti-human IgG Fc Capture (AHC) Biosensors (Sartorius). After reaching a displacement of 0.8–1 nm, biosensors were transferred into BLI buffer for a second baseline. For association step, biosensors were moved into wells containing serial dilution of CD80-His, and finally transferred into wells containing BLI buffer for dissociation step. Kinetic results were fit to a 1:1 model using global fitting, with y-axis alignment on the average of a segment of the dissociation step, using the Octet® Analysis Studio Software.

Avidity measurements were performed as above, with CD80-His immobilised with Nickel NTA-tips (Sartorius). For association step, serial dilutions of CTLA-4 constructs were made at 50, 25, 12.5, 6.25, 3.125, 1.56, 0.78 nM.

### CTLA-4 and PD-1 cell binding assays

PD-1 Ig (Bio-Techne), CTLA-4 Ig (Abatacept) and in-house generated CTLA-4 and CD28 Ig constructs were conjugated with APC or Dy405 using the Lightning-Link® Conjugation kits (Abcam), according to manufacturer’s instructions. DG-75 cells were incubated with indicated concentration of conjugated CTLA-4 or CD28 constructs for 30 mins at 37°C. Cells were washed once in PBS followed by 15 mins stain with 1 μg/ml of PD-1 Ig – Dy405, at 37°C. Cells were washed twice in PBS followed by fixation in 4% PFA.

For *trans*-endocytosis assays, DG-75 cells were first stained with Cell Trace Violet (ThermoFisher Scientific) according to manufacturer’s instructions. 1 × 10^5^ DG-75 cells were co-cultured with 1 × 10^5^ Jurkat T-cells expressing either WT CTLA-4, CTLA-4-Del36 or no CTLA-4. Wells were supplemented with 5 ng/ml of Staphylococcal Enterotoxin E (SEE, Biomatik) to enhance cell-cell contacts. Assay was carried out in round-bottom, 96-well plate, in a total volume of 200 μl, for 24 hrs at 37°C. Cells were then washed once in PBS followed by 15 mins stain with 1 μg/ml of PD-1 Ig – APC, at 37°C. Cells were washed twice in PBS followed by fixation in 4% PFA. In all cases, cells were resuspended in FACs buffer and acquired on the BD Fortessa.

### Statistical and *in silico* analysis

Statistical analyses and significance were determined using GraphPad Prism v9.02 software (GraphPad Software Inc.). Experiments were performed in triplicate and statistical significance determined with appropriate test, as indicated in figures.

Flow cytometry data was post-processed with FlowJo (BD Biosciences). Structural models of proteins were accessed from PDB and analysed with UCSF ChimeraX. Alignment was performed using the matchmaker tool.

## Acknowledgements

AK, EW, CH, and DMS were funded by the Wellcome Trust (Grants 204798/Z/16/Z). MAR and LF were funded by an MRC industrial collaborative studentship with Astra Zeneca. DG was funded by an MRC-DTP studentship. This research was funded in whole, or in part, by the Wellcome Trust (Grant 204798/Z/16/Z).

For the purpose of Open Access, the author has applied a CC BY public copyright licence to any Author Accepted Manuscript version arising from this submission.

---

**Fig S1.**
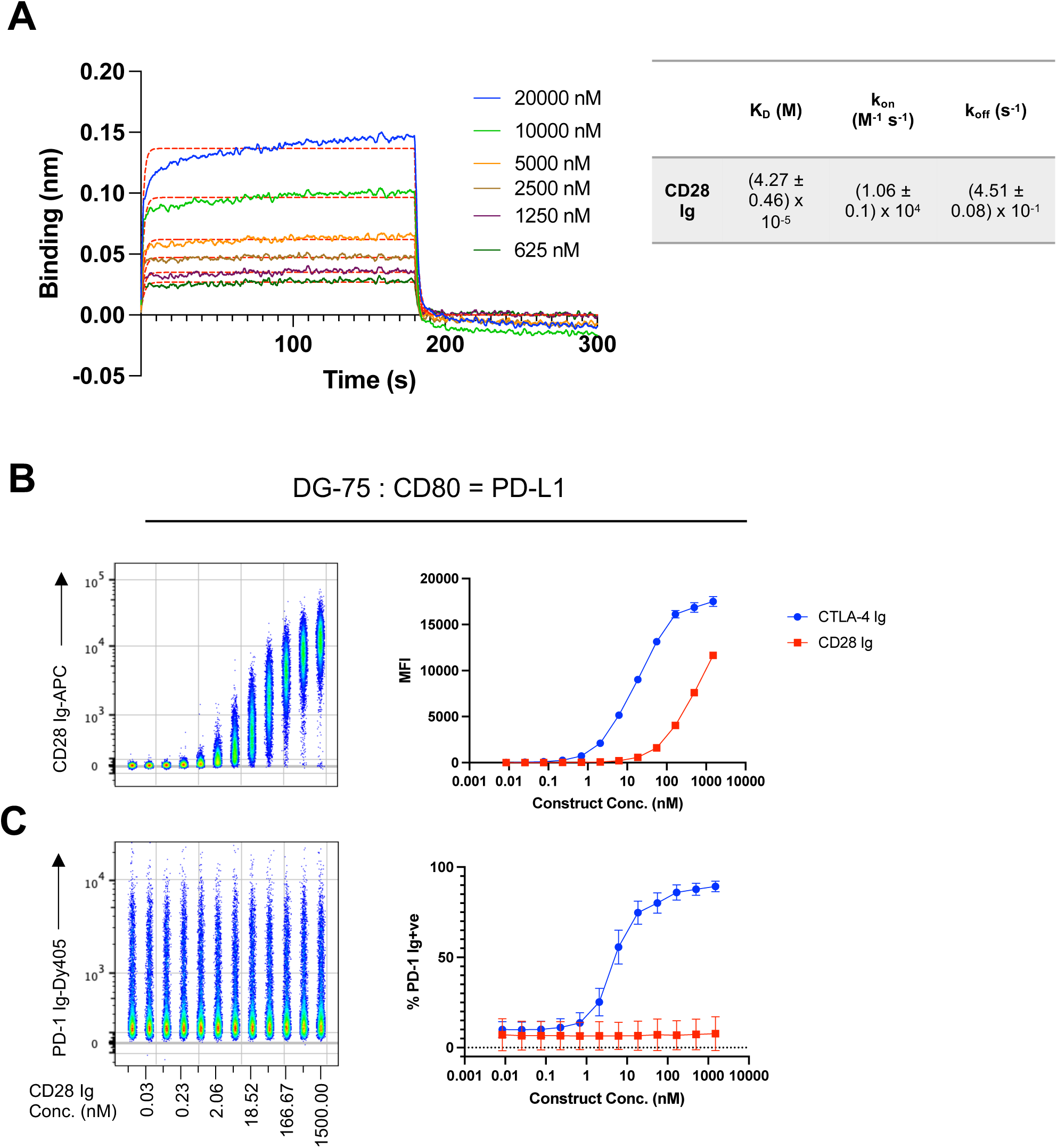
Soluble CD28 Ig fails to disrupt the CD80 PD-L1 interaction. (A) Binding curves showing the association and dissociation of 20, 10, 5, 2.5, 1.25, 0.625 uM of CD80-His to CD28-Ig. Red lines show best fit to a 1:1 binding model. Right hand table details kinetic and thermodynamic parameters (K_D_, k_on_, k_off_) obtained from the best global fit of the association/dissociation data to a 1:1 binding model. The errors given are fitting errors from the global fitting. **(B)** Concatenated flow cytometry plot of a 12-point serial dilution of CD28 Ig-APC, starting at 1500nM, on DG-75 : CD80 = PD-L1, with graphical representation in right-hand panel. **(C)** PD-1 Ig binding on cells described in (B), with graphical representation in right-hand panel. Data are representative of three independent experiments showing mean ± SD.

**Fig S2.**
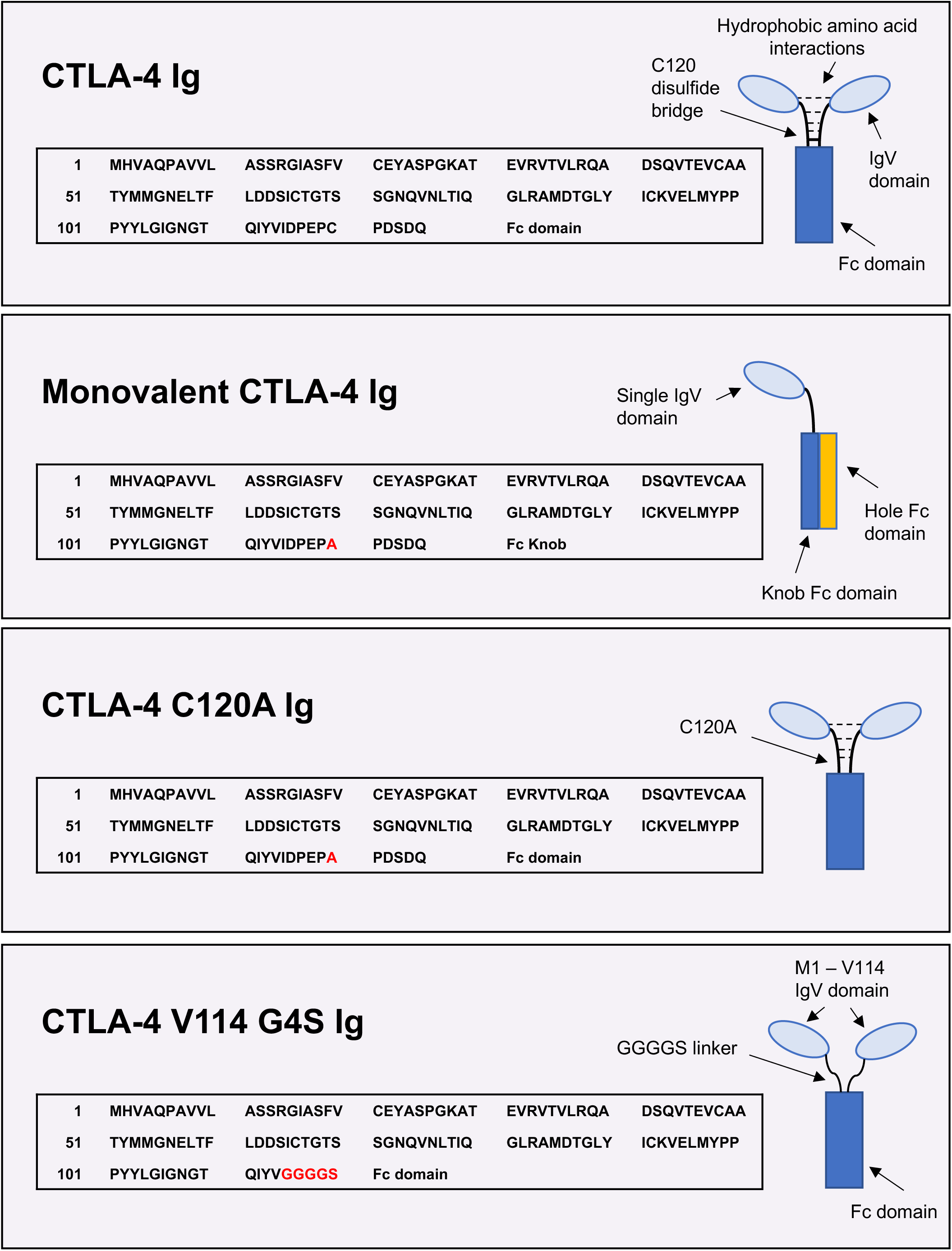
Amino acid sequences and schematics of CTLA-4 – Fc constructs.

**Fig S3.**
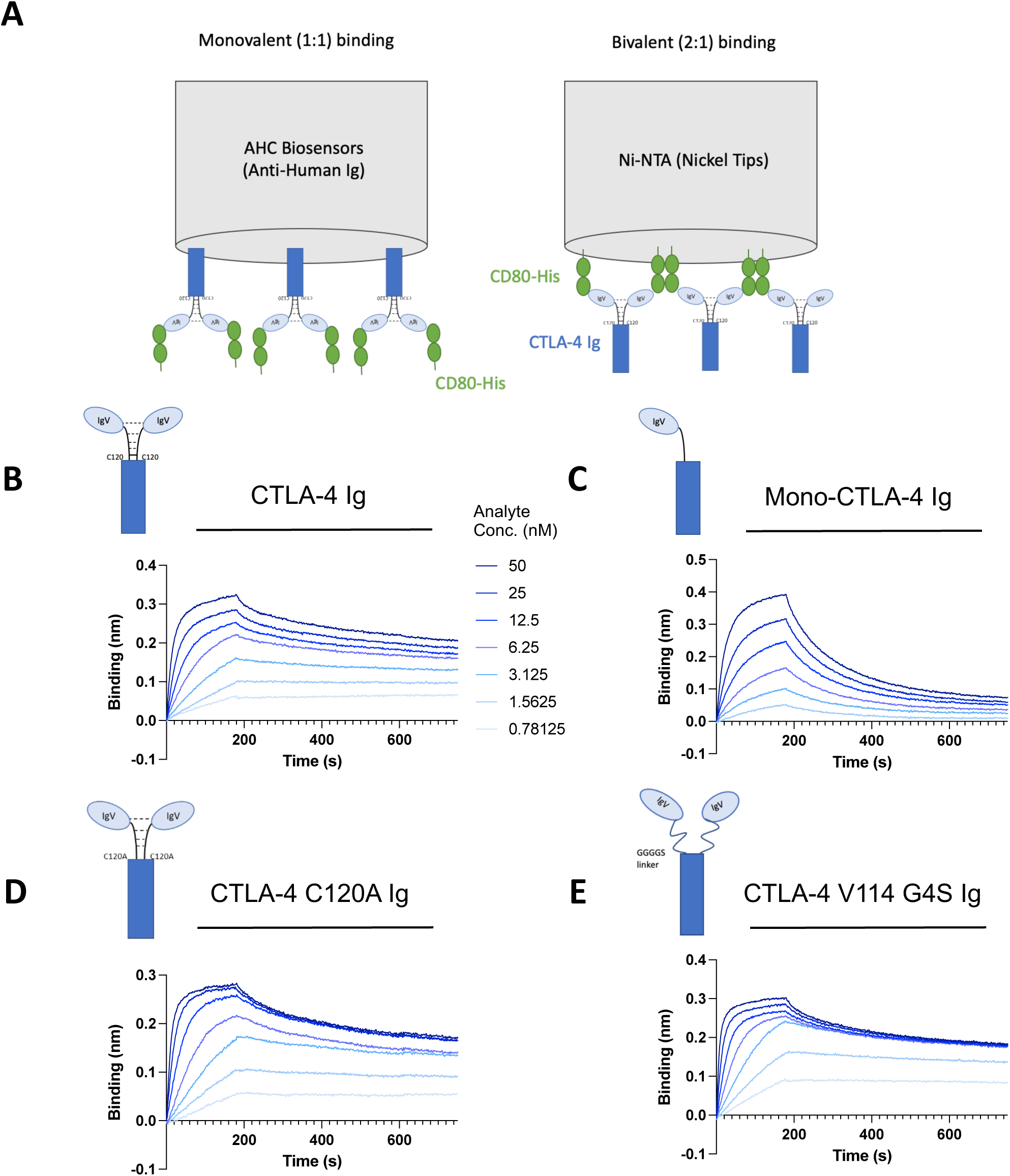
All bivalent CTLA-4 constructs bind CD80 with enhanced avidity. **(A)** Schematic of Bio-Layer Interferometry assays assessing 1:1 and 2:1 binding models. **(B-D)** Binding curves showing the association and dissociation of 50, 25, 12.5, 6.25, 3.125, 1.5625, 0.78125 nM of indicated CTLA-4 Ig constructs to CD80-His.

**Fig S4.**
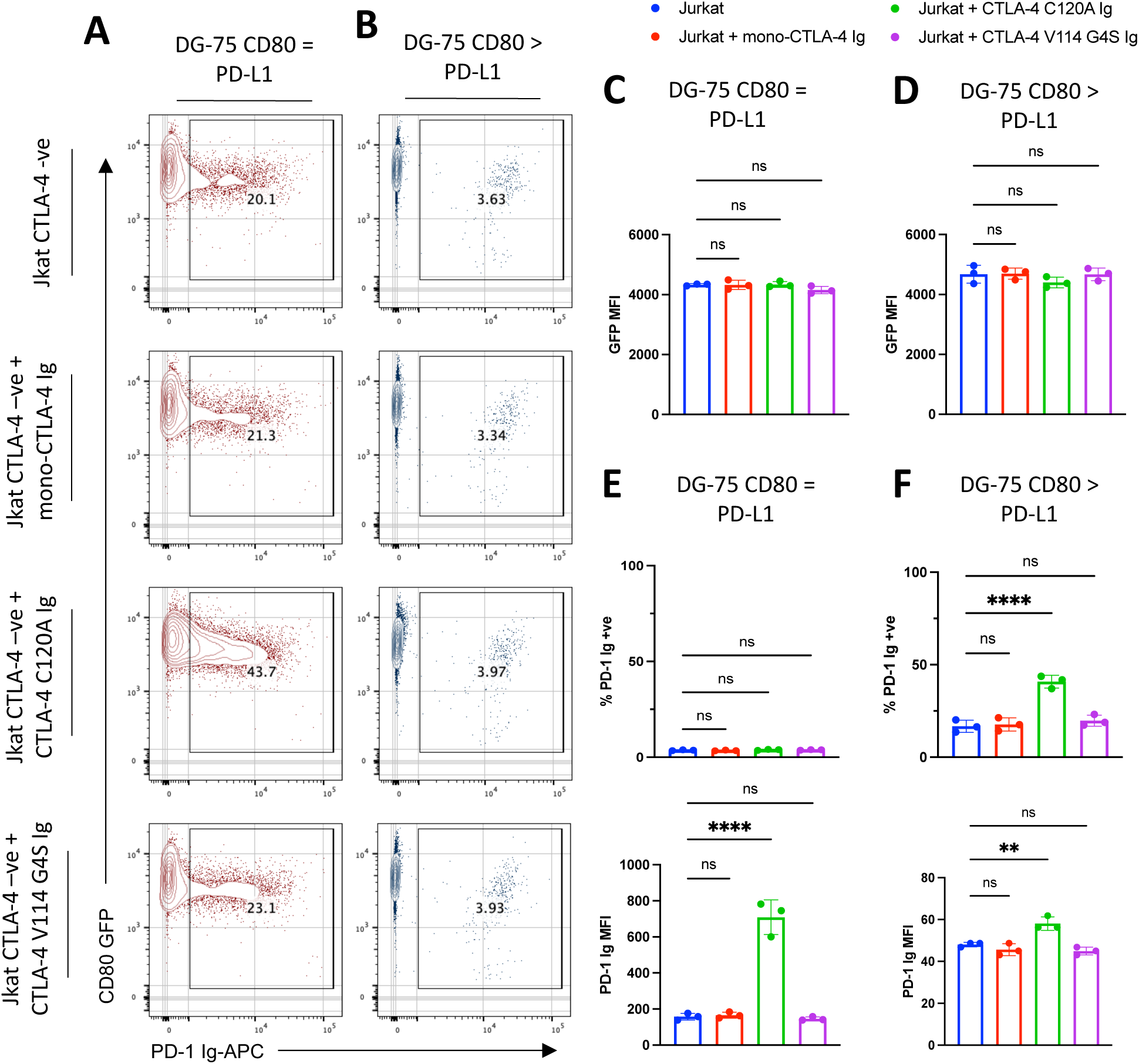
Monovalent and flexible soluble CTLA-4 constructs fail to restore PD-1 binding. **(A & B)** DG-75 : CD80 = PD-L1 (A) and DG-75 : CD80 > PD-L1 (B) incubated for 24hrs with either: CTLA-4-ve Jurkat, CTLA-4-ve Jurkat + 50nM mono-CTLA-4 Ig, CTLA-4-ve Jurkat + 50nM CTLA-4 V114 G4S Ig and CTLA-4-ve Jurkat + 50nM CTLA-4 C120A Ig. Cells were stained with 1ug/ml of PD-1 Ig-APC after incubation. Data shows representative FACS plots of CD80-GFP vs. PD-1 Ig. **(C-F)** Graphical representation of (A & B) respectively, plotting CD80-GFP MFI, % of PD-1 Ig +ve cells and PD-1 Ig MFI on DG-75 : CD80 = PD-L1 (C,E) and DG-75 : CD80 > PD-L1 (D,F). Data are representative of three independent experiments showing mean ± SD. **P ≤ 0.01, ****P ≤ 0.0001, ns, not significant: one-way ANOVA with Tukey’s multiple comparisons test (C-F).

